# Identification of a volatile chemotype associated with resilience to water stress in domesticated varieties of Cotton

**DOI:** 10.1101/2025.07.16.665191

**Authors:** Christopher J. Frost, Sarah M. Johnson, Duke Pauli, Giovanni Melandri

**Affiliations:** The BIO5 Institute, University of Arizona, Tucson, AZ 85719; School of Plant Sciences, University of Arizona, Tucson, AZ 85721; Center for Agroecosystem Research in the Desert (ARID), University of Arizona, Tucson, AZ 85721

## Abstract

Volatile organic compounds (VOCs) mediate plant responses to environmental stresses, yet their chemotypic variation and drought responsiveness remain poorly characterized in domesticated crops. Here, we identify a previously undocumented binary chemotype structure in cultivated upland cotton (*Gossypium hirsutum*), defined by the mutually exclusive production of either bisabolene or guaiene sesquiterpenes. Across 24 genotypes grown under field conditions, genotypes consistently produced one of these compounds, but never both, revealing a cryptic chemotypic polymorphism with physiological consequences. While drought stress reduced leaf water content across all plants, plants in the guaiene chemotype exhibited disproportionately greater losses. Water limitation also reduced total monoterpene and sesquiterpene concentrations, with the bisabolene chemotype showing stronger declines despite higher constitutive levels. Chemotypes further diverged in their biosynthesis of key damage-induced green leaf volatiles (GLVs). While the guaiene chemotype reduced GLV biosynthetic capacity under prolonged drought, the bisabolene chemotype showed transient increases in GLVs and pathway-specific enzyme activity under short-term stress, suggesting differential physiological resilience. Notably, all genotypes in this study fell within the low γ-terpinene category previously described in wild cotton, indicating that this bisabolene/guaiene variation represents a novel axis of chemical diversity distinct from known wild chemotypes. These findings position chemotypic variation as a promising, non-destructive biomarker for drought response in cotton breeding programs. By linking chemotype to constitutive VOCs and water status, our work provides new tools for identifying stress-resilient cultivars and advances understanding of phytochemical strategies for crop management in a changing climate.

## Introduction

Plants synthesize and emit volatile organic compounds (VOCs), a diverse group of secondary metabolites that mediate responses to a wide range of environmental stresses. While VOCs are well known for their roles in plant–insect and plant–pathogen interactions ^1–11^, their roles in abiotic stress responses ^12^, particularly under drought conditions ^13,14^, are increasingly recognized as critical for plant acclimation and survival during dynamic growing seasons.

VOCs differ in their biosynthetic origins, persistence, and ecological function. Terpenes, one of the most common classes of VOCs, may be synthesized de novo in response to stress or stored constitutively in specialized structures such as glandular trichomes or secretory cavities. In species that store terpenes, low levels are often passively released under normal conditions, but such pre-storage allows for large quantities to be emitted rapidly when tissues are damaged. Another important class of plant VOCs are the so-called green leaf volatiles (GLVs), six-carbon aldehydes, alcohols, and acetates derived from linolenic acid via the lipoxygenase pathway ^15^, are almost exclusively synthesized de novo following mechanical damage ^16^, herbivory ^6^, or abiotic stress that causes tissue disruption. GLVs serve several adaptive functions, including a role as infochemicals due to their ability to trigger defensive or acclimatory responses in neighboring plants and attract or repel herbivores and their natural enemies ^17^.

VOC profiles can vary both within and between plant species, and substantial intraspecific variation has been documented in wild and domesticated taxa. Several wild species exhibit genetically based “chemotypes,” or discrete chemical polymorphisms defined by dominant terpenes ^18^. These chemotypes influence not only plant interactions with herbivores and pathogens ^19,20^, but also susceptibility to abiotic stress ^21^ and capacity for between-plant signaling ^22^. Domestication can further alter VOC production, leading to differences in constitutive and inducible emissions between cultivated and wild populations ^23–26^. Because secondary metabolites are often reduced during crop domestication ^23,24^, identifying chemotype-linked VOC traits in modern varieties presents an opportunity to develop non-destructive biomarkers for breeding stress-resilient crops.

Upland cotton (*Gossypium hirsutum* L.) holds substantial agricultural and economic importance within the United States, contributing significantly to the nation’s fiber and agricultural sectors. Cotton’s economic footprint extends well beyond initial farm sales, supporting numerous jobs and contributing billions to the economy via exports and related industries ^27^. Arizona, for instance, consistently ranks among the top cotton-producing states nationally and exceeding $32M in cottonseed production in 2024 ^28^. However, the sustainability of cotton production, especially in arid and semi-arid regions such as Arizona where this experiment was conducted, is increasingly challenged by the escalating frequency and intensity of abiotic stresses ^29^. Drought is a primary abiotic environmental constraint that severely impedes cotton yield potential and production efficiency worldwide ^30^. Identifying novel strategies to evaluate drought resilience could significantly support and accelerate breeding efforts aimed at developing new drought tolerant cotton varieties.

Cotton plants constitutively produce and store mono- and sesquiterpenes in foliar glands, which enables large, instantaneous VOC release upon tissue disruption by herbivores or mechanical damage ^31^. Emissions from damaged cotton leaves typically comprise a mixture of stored terpenes and wound-induced GLVs, which are biosynthesized within second of wounding ^16^. While VOCs have been extensively studied in the context of herbivory ^32–34^, relatively little is known about how terpene and GLV emissions vary among domesticated cotton genotypes or respond to abiotic stress ^35^ or whether VOCs represent viable biomarkers. In wild *G. hirsutum* populations from the Yucatán Peninsula, two heritable chemotypes, distinguished by high or low γ-terpinene, have recently been identified ^36^. However, domesticated cotton varieties are almost exclusively low γ-terpinene producers ^34^, limiting the functionality of this chemotype as a biomarker of environmental responses in domesticated varieties. Given the narrow genetic base of modern cotton cultivars ^37^, identifying new chemotypic variation in VOC production could provide critical tools for improving resilience to environmental stressors.

Drought stress profoundly influences plant metabolism, notably affecting the biosynthesis and emission of VOCs ^38,39^. Plants often exhibit a reduction in the emission of constitutive ^40^ and inducible ^41^ terpenes, including both monoterpenes and sesquiterpenes, due largely to constrained carbon assimilation and limitations in substrate availability. Drought-mediated reductions in terpene biosynthesis could, in turn, lead to heightened susceptibility to biotic stress since these volatile terpenes influence herbivores and pathogens ^42^. Conversely, the GLV biosynthesis capacity and emissions may actually increase under drought stress ^43^, potentially enhancing some aspects of plant resilience by priming defense mechanisms ^2,4,5^ or by attracting beneficial organisms ^44,45^. Relatively little is known about how terpene and GLV emissions vary among domesticated cotton genotypes or respond to drought stress ^35^. Recent work with hyperspectral analysis in upland cotton under drought and heat stress has revealed the important linkage between drought stress and the overall cotton phytochemical metabolome ^29^, suggesting that phytochemical markers could play a role in predicting drought resistance. Therefore, understanding these responses is crucial, as they may inform the development of non-destructive physiochemical biomarkers for breeding drought-resilient cotton varieties and other crops where non-destructive, early indicators of physiological performance are urgently needed.

Here, we characterize a previously undocumented cryptic chemotype system in domesticated cotton, defined by the exclusive presence of either bisabolene or guaiene sesquiterpenes. While analyzing in vitro VOC emissions from 24 upland cotton genotypes grown in Arizona under field conditions, we observed that each genotype produced either bisabolene or guaiene isomers, but never both. Of the 24 genotypes studied, 17 were classified within the bisabolene-producing chemotype, 6 within the guaiene-producing chemotype, and one (TM-1) as unclassified in that it produced neither bisabolene nor guaiene under the conditions tested. We evaluated the influence of chemotype and water availability on in vitro biosynthesis of monoterpenes, sesquiterpenes, and GLVs. By linking VOC chemistry to chemotypic and genotypic variation, we highlight the potential for using chemotypic variation as a non-invasive biomarker to inform next-generation cotton breeding for drought resilience and environmental sustainability while addressing the need for more sustainable cultivation practices, particularly in arid and drought-prone environments.

## Methods

### Field experiment

Twenty-four cotton (*Gossypium hirsutum*) accessions (Supp. Table S1) were grown during the summer of 2023 at the University of Arizona’s Maricopa Agricultural Center (33°04ʹ37ʺN, 111°58ʹ26ʺW) in central Arizona. The experiment followed a randomized incomplete block design with four replicated plots per genotype, totaling 96 experimental plots. Plants were irrigated using a drip tape system placed at a depth of 20 cm. Following crop establishment and early canopy development, a water-limitation (WL) treatment was applied to half of the plots, while the remaining plots were maintained under well-watered (WW) conditions. In other words, the four plots were split equally among the treatments with two biological replicates being subjected to each water treatment. Differential irrigation treatments were maintained for the remainder of the field season. Soil volumetric water content (SVWC%) was monitored throughout the growing season using a field-calibrated neutron probe (model 503; Campbell Pacific Nuclear, CPN, Martinez, CA) with 16 neutron access tubes installed throughout the field (Supp. Figure S1). Measurements were taken at three depths (30, 50, and 70cm) to capture soil moisture dynamics across the root zone.

Leaf tissue was collected for in vitro VOC analysis at two separate time points during flower development, July 21st and August 3rd. These dates are hereafter referred to as Time Point 1 (TP1) and Time Point 2 (TP2). This sampling regime was intended to capture physiological status at three weeks (TP1) and five weeks (TP2) after the onset of chronic water limitation, representing relatively short-term and longer-term drought conditions and roughly corresponding to ‘early’ and ‘late’ stage of flowering for the cotton genotypes. Sampling occurred between 10:00am and 12:30pm. Leaf punches from five representative plants per plot were collected, immediately snap frozen in liquid nitrogen, and stored for VOC analysis. From the same leaves, additional leaf punches were collected for determination of leaf water content.

### VOC Collections

We collected VOCs using vapor phase extraction directly from cryogenic leaf material. Cryogenic leaf material was ground into a fine powder using a ball mill grinder (Retsch MM300, Germany), and 10 mg fresh weight (FW) per sample of this material was weighed into a new vial. Under cryogenic conditions (liquid nitrogen), the vial was connected to a Teflon tube connected to a vacuum flow meter with a VOC collection filter placed in line. Theses VOC collection filters are routinely used for VOC sampling ^46^, and each filter contained ∼30 mg of Poropak Q. The flow meter was set to 400 ml / min and the sample was removed from the cryogenic conditions to thaw to room temperature, allowing for the release of volatile terpenes and enabling the biosynthesis of wound-related GLVs ^16^. VOC collections occurred for 60 min.

Preliminary testing established this method as a robust approach to sampling pre-synthesized terpenes and rapidly synthesized GLVs without additional de novo synthesis of terpenes (Supplemental Fig. S2). This method is similar to hexane extractions of volatile terpenes as has been performed previously with cotton ^36,47^, but benefits from avoiding the contamination of pigments and other hexane-soluble elements being injected onto the gas chromatograph and avoids the need for solid-phase extraction.

VOCs collected in the filters were then eluted with 100 µl of dichloromethane containing 5 ng / µl of tetralin as an internal standard directly into a microinsert seated inside GC autosampler vial.

### GC-MS Profiling

VOCs were resolved and quantified on an Agilent 8890 Gas Chromatograph (GC) coupled with a 7000D Triple Quadrupole Mass Spectrometer (MS) operating in scan mode. One μl of each sample was injected into the GC in splitless mode on a single taper Ultra Inert liner (Agilent #5190-2293) with inlet temperature and transfer line set to 250°C. VOCs were resolved on a DB-5MS column (30 m length, 0.25 mm diameter, 0.25 μM film thickness, 10 m Duraguard pre-column; Agilent #122-5532G) with a constant flow of 1.0 ml/min, and an average velocity of 21.98 cm/sec. Thermal ramping initiated at 35°C for 1 min, ramped at 20°C / min to 250°C, and held for 10 min. The MS operated with an EI ion source (70 eV) with the MS in scanning mode (40-400 m/z). The transfer line, the ion source, and the quadrupole temperatures were set to 250°C, 230°C, and 150°C, respectively.

We used MassHunter Quantitative (Agilent, version 10.2) software to quantity the VOCs. Individual VOCs were quantified based on a single *m/z* ion, with at least one qualifying *m/z* ion ( Supplemental Table S2). Peak areas were normalized against the internal standard (tetralin) and quantified against external standard curves (Supplemental Fig. S3; Supplemental Table S3). In cases where an authentic standard was not available, this VOC was quantified against the most structurally similar VOC (Supplemental Table S2).

### Statistical Analysis

VOC concentrations were analyzed using linear mixed-effects models (LMMs) implemented via the *lmer* function in the *lme4* package ^48^ in R (version 4.2.1) ^49^. For chemotype-level analyses, fixed effects included ‘water treatment’, ‘time point’, and ‘chemotype’, while random effects accounted for ‘genotype’, ‘plot’, and ‘replicate’. Each model was evaluated for assumptions of normality and homoscedasticity, and data were log- or square root-transformed as needed to meet model assumptions. To assess genotype-specific effects, a parallel model structure was used, with ‘genotype’, ‘water treatment’, and ‘time point’as fixed effects, and ‘plot’ and ‘replicate’ as random effects.

Analysis of variance (ANOVA) was conducted using the *Anova* function in the *car* package ^50^, and post hoc pairwise comparisons were performed using the *emmeans* package ^51^ with Tukey-adjusted p-values. Compact letter displays (CLDs) were generated using the *cld* function in the *multcomp* package ^52^ to support visual interpretation of significant group differences. All visualizations were generated using the *ggplot2* package ^53^.

## Results

### Determination of Chemotypes and Effects on Leaf Water Content

VOC profiles revealed two closely eluting peaks at a retention time of approximately 13 minutes, separated by only 0.02 min, making them easily overlooked in chromatographic data (Fig. 1A, B). In the absence of mass spectral information, they would easily be passed over as identical compounds. However, mass spectral analysis confirmed that these peaks correspond to distinct compounds, putatively identified as y-guaiene and bisabolene (Fig. 1C). Spectral matching against the NIST14 library was 95.7 +/- 1.3 % and 91.4 +/- 1.6 % for y-guaiene (hereafter ‘guaiene’) and bisabolene, respectively. Notably, the presence of these compounds was entirely genotype-dependent; genotypes that produced one compound were consistently devoid of the other, indicating a mutually exclusive chemotypic pattern (Supplemental Fig. S4). As further confirmation of chemotypic distinction, the guiaene chemotype also produced α-guaiene (RT ∼ 12.357 m), which was unique to that chemotype and not produced in the bisabolene chemotype.

**Figure 1.**
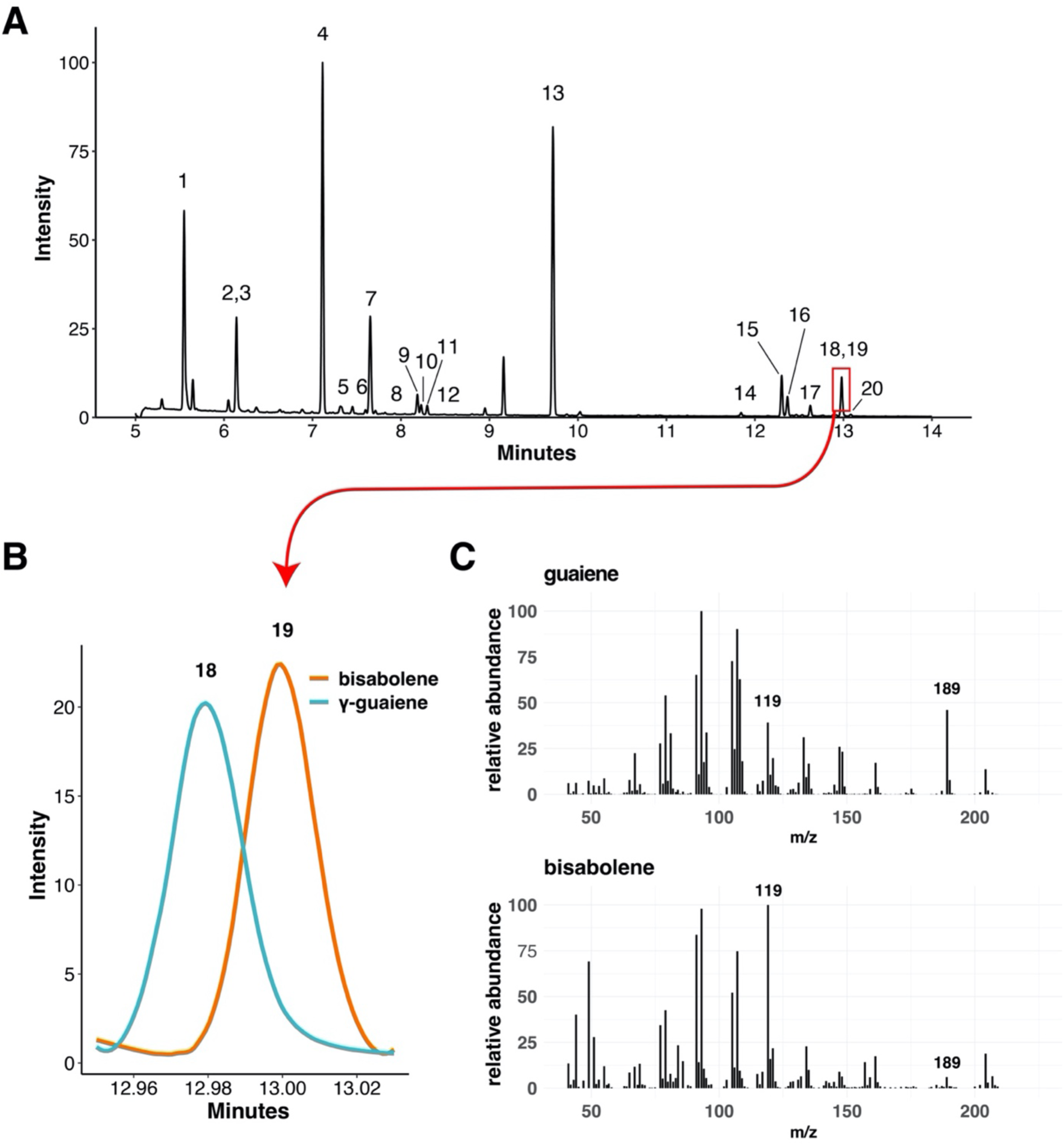
Total ion chromatogram of cotton in vitro VOCs and library matches to bisabolene and γ-guaiene. (A) Representative total ion chromatogram. 1: (*Z)*-3-Hexenal; 2: (*E*)-2-Hexenal; 3: (*Z*)-3-Hexen-1-ol; 4: α-Pinene; 5: Camphene; 6: β-Myrcene; 7: β-Pinene; 8: α-Phellandrene; 9: D-Limonene; 10: β-Phellandrene; 11: β-Ocimene; 12: γ-Terpinene; 13: Tetralin (internal standard); 14: α-Copaene; 15: β-Caryophyllene; 16: α-guaiene; 17: Humulene; 18: γ-guaiene; 19: bisabolene; 20: γ-Cadinene. (B) Total ion chromatogram showing elution of bisabolene and γ-guaiene. (C) Mass spectra of corresponding bisabolene and guaiene peaks. NIST library matches for γ-guaiene and bisabolene were scored on average (across all genotypes) at 95.7 +/- 1.3 % and 91.4 +/- 1.6 %, respectively.

As expected, samples collected from water limited (WL) plots displayed a severely reduced leaf water content (LWC%) compared with the ones collected from well-watered (WW) plots (Fig. 2A). However, there was also a chemotype effect on leaf water content, where the the guaiene chemotype experienced a greater proportional reduction in leaf water content relative to the bisabolene chemotype (Fig. 2B).

**Figure 2.**
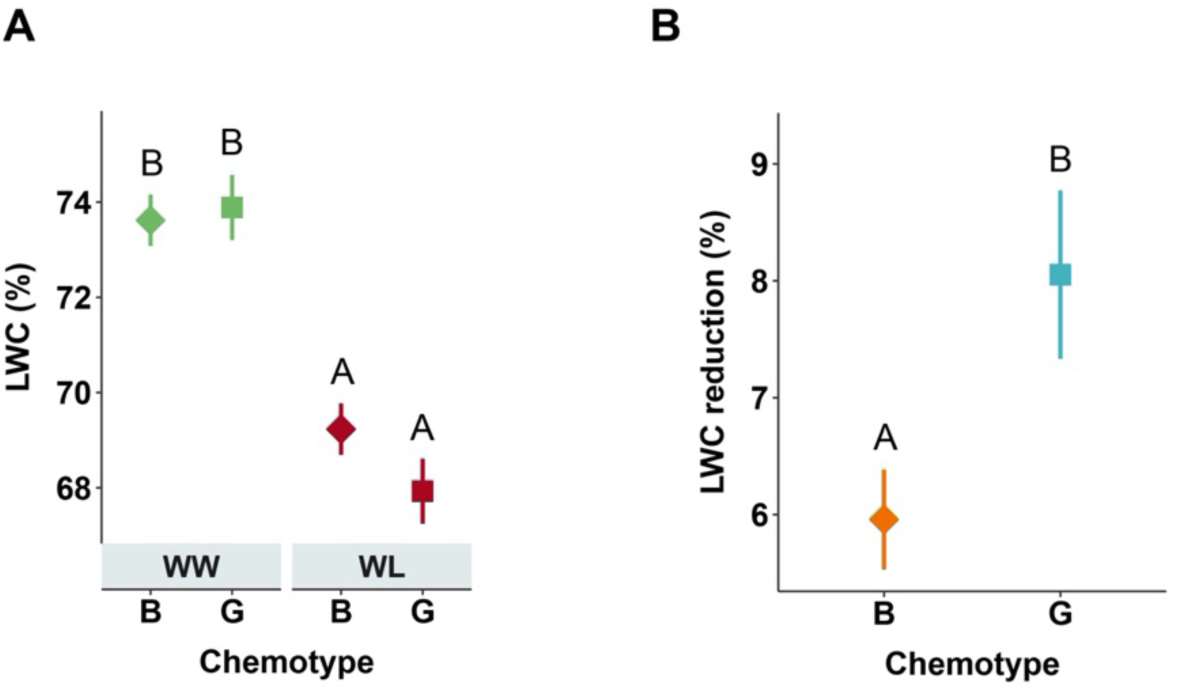
Leaf water content response to water limitation varies by chemotype. (A) Mean leaf water content (LWC%) of leaves sampled for volatile analysis under well-watered (WW; green symbols) and water-limited (WL; red symbols) conditions for two chemotypes, bisabolene (B) and guaiene (G). LWC% calculated as the proportion of water lost after drying relative to the fresh weight. (B) Leaf water loss under drought, calculated as the difference in LWC% between water-limited and well-watered treatments relative to the well-watered treatment in two chemotypes. Symbols represent means ± SE. Letters denote significant differences among groups (*p* < 0.05) based on Tukey-adjusted pairwise post hoc comparisons performed using the *emmeans* package in R.

### Chemotype-Specific Monoterpene Responses to Water Limitation

Total monoterpene concentrations were strongly reduced under water-limited conditions, regardless of chemotype or the duration of the water limitation treatment (Fig. 3A). There was also a strong main effect of chemotype, wherein the bisabolene chemotype produced more monoterpenes overall than did the guaiene chemotype. This trend was evident for α-pinene, the most abundant monoterpene, as well as several less abundant compounds, including limonene and β-phellandrene (Fig. 3B). However, several individual monoterpenes exhibited chemotype-specific responses to water limitation. For example, β-pinene and β-ocimene concentrations were higher in the bisabolene chemotype compared to the guaiene chemotype under short-term water limitation (TP1), but these chemotypic differences were not maintained under longer-term water limitation (TP2) (Fig. 2B). Camphene was the only compound to show chemotypic variation across both time points; its concentrations were consistently low in the guaiene group, resulting in no detectable reduction under water limitation. In contrast, γ-terpinene, previously highlighted in chemotype-level studies of wild cotton ^36^, was reduced under water limitation in the bisabolene chemotype, with no corresponding change in the guaiene group. Lastly, β-myrcene and α-phellandrene exhibited minimal variation across treatments or chemotypes.

**Figure 3.**
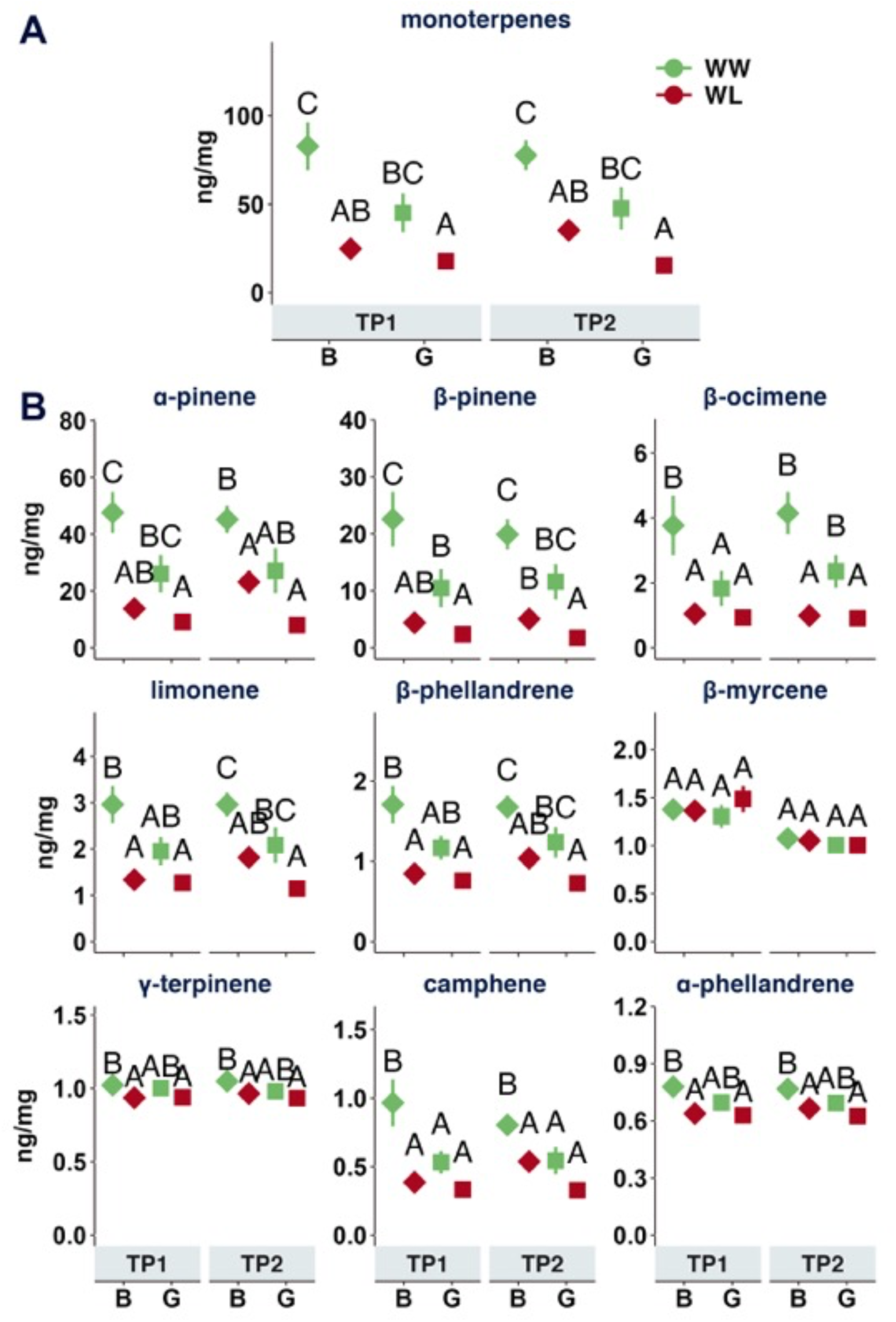
**Chemotype-specific responses of monoterpenes to water limitation in domesticated cotton**. (A) Total monoterpene concentrations (ng/mg) measured in leaf tissue under well-watered (WW; green symbols) and water-limited (WL; red symbols) conditions for two chemotypes, bisabolene (B) and guaiene (G), at two time points. TP1 represents short-term water limitation and TP2 represented long-term water limitation. (B) Concentrations (ng/mg) of individual monoterpenes across treatments, chemotypes, and time points. Symbols represent treatment means ± SE. Letters denote significant differences among groups (*p* < 0.05) based on Tukey post hoc comparisons performed using the *emmeans* package in R.

### Effects of Water Limitation on Sesquiterpene Concentrations Vary by Chemotype

The bisabolene chemotype exhibited consistent reductions in total sesquiterpene concentrations under both short-term (TP1) and longer-term (TP2) water limitation, a pattern not observed in the guaiene chemotype (Fig. 4A). This stability in the guaiene chemotype appeared to be due to relatively low constitutive sesquiterpene concentrations under well-watered conditions. Among the four sesquiterpenes included in the total calculation, β-caryophyllene, humulene, and α-copaene followed the overall reduction pattern in the bisabolene chemotype, while γ-cadinene remained low and stable across chemotypes and treatments (Fig. 4B). bisabolene and guaiene compounds were excluded from total sesquiterpene calculations because they define the respective chemotypes and could disproportionately influence total concentrations. Nevertheless, in the bisabolene chemotype, bisabolene concentrations declined at both TP1 and TP2, and α-bisabolene epoxide was reduced under long-term water limitation (Fig. 4C). In the guaiene chemotype, γ-guaiene and α-guaiene concentrations were lower under long-term water limitation relative to well-watered controls (Fig. 4C).

**Figure 4.**
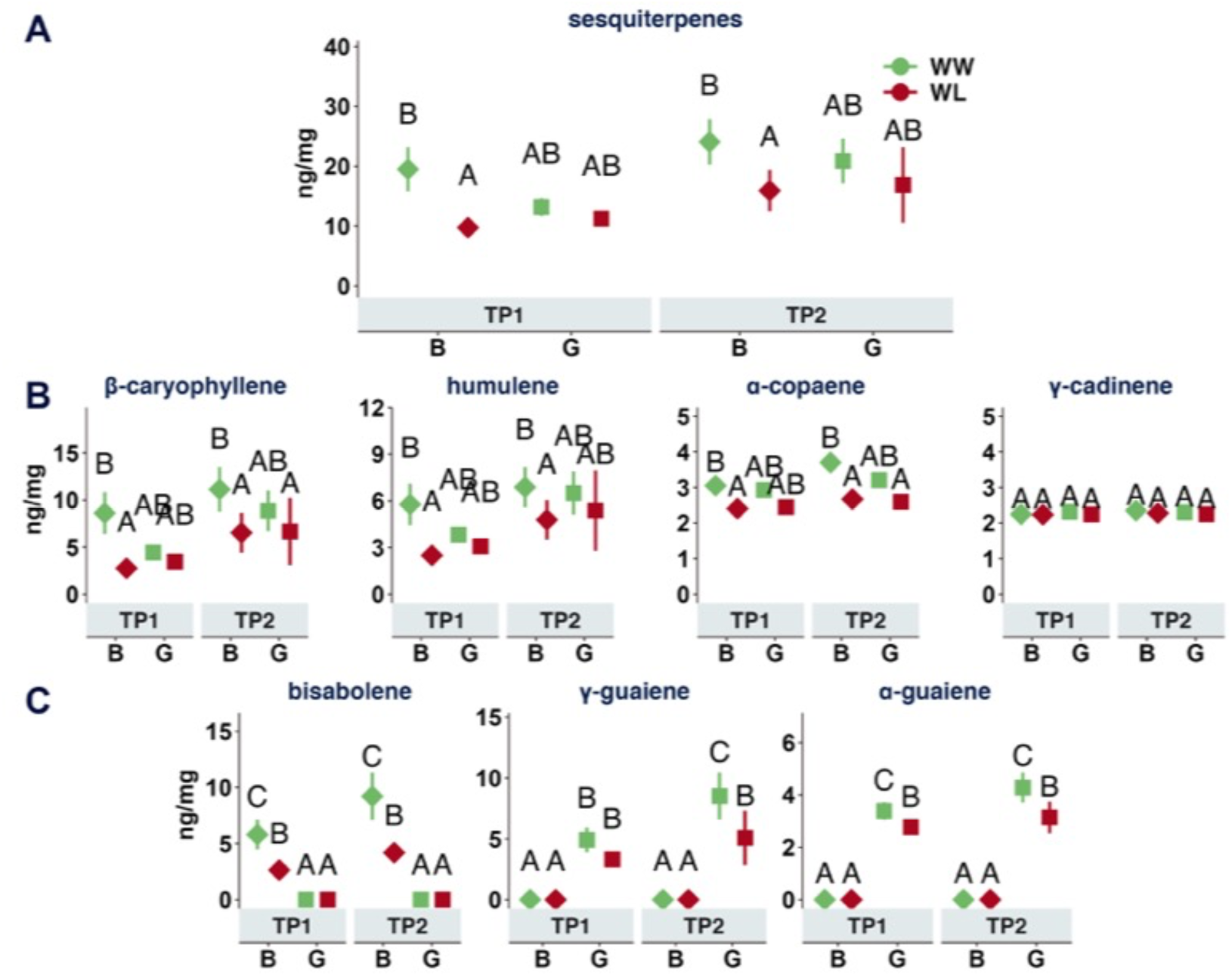
Chemotype-specific responses of sesquiterpenes to water limitation in domesticated cotton. (A) Total sesquiterpene concentrations (ng/mg) measured in leaf tissue under well-watered (WW; green symbols) and water-limited (WL; red symbols) conditions for two chemotypes—bisabolene (B) and guaiene (G)—at two time points: TP1 (short-term water limitation) and TP2 (long-term water limitation). bisabolene- and guaiene-defining compounds were excluded from total concentrations to avoid chemotype-driven bias. (B) Concentrations (ng/mg) of individual sesquiterpenes included in the total calculation: β-caryophyllene, humulene, α-copaene, and γ-cadinene. (C) Concentrations of the chemotype-defining sesquiterpenes: bisabolene, γ-guaiene, and α-guaiene. Symbols represent treatment means ± SE. Letters denote significant differences among groups (*p* < 0.05) based on Tukey-adjusted pairwise post hoc comparisons performed using the *emmeans* package in R.

### GLV Biosynthesis Capacity varies by Chemotype and Duration of Water Limitation

GLVs are key VOCs emitted in response to physical damage and various abiotic stresses, including drought. As infochemicals, GLVs play important ecological roles such as mediating plant-to-plant signaling. The three primary damage-induced GLVs are synthesized de novo from linolenic acid via a well-characterized pathway involving four enzymes (Fig. 5A). Because GLV biosynthesis is activated within seconds of tissue damage and is not pre-stored ^16^, the in vitro GLV measurements reported here reflect the constitutive biosynthetic capacity and activity of this enzymatic cascade under field conditions.

**Figure 5.**
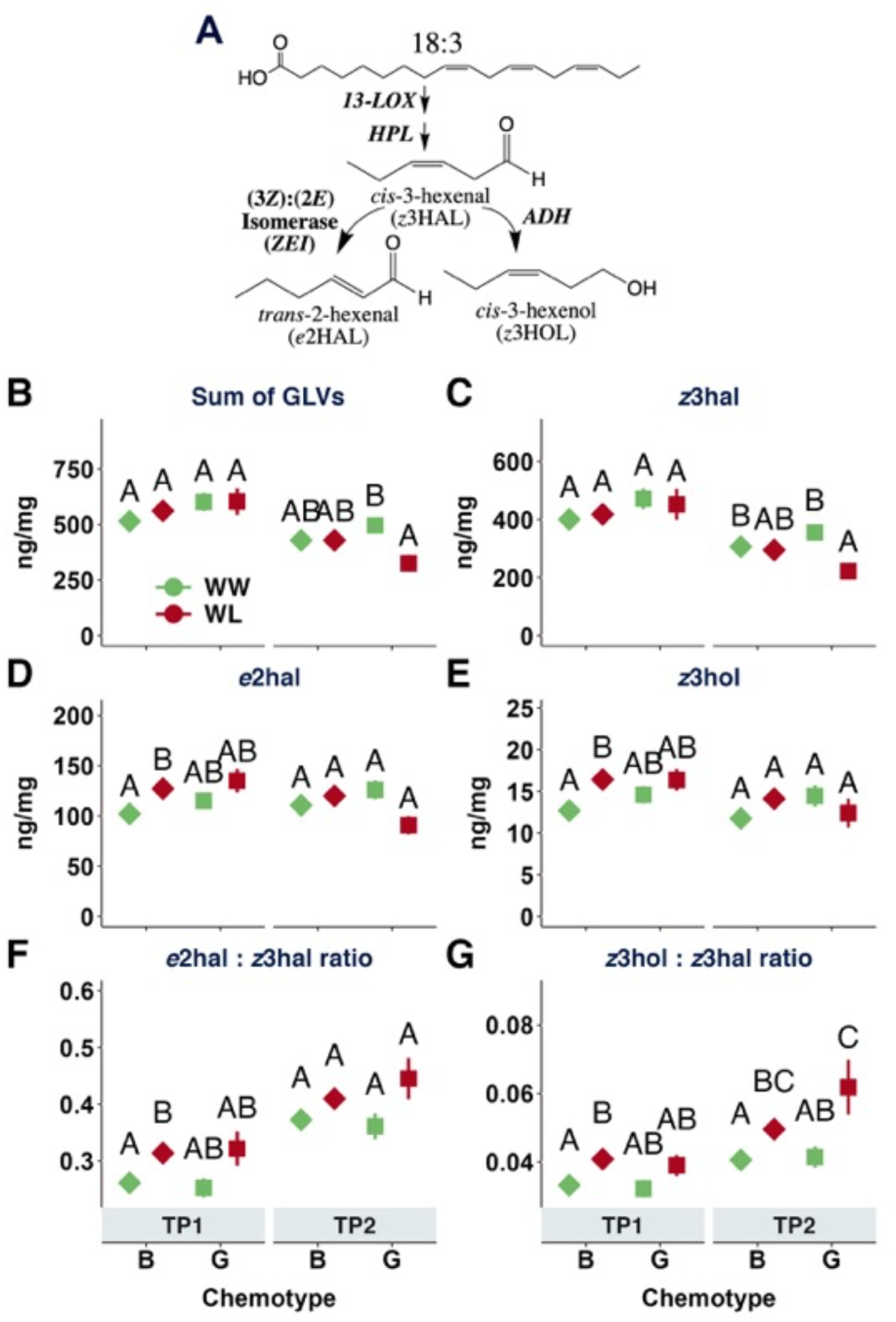
Chemotype-specific responses of green leaf volatile (GLV) biosynthesis to water limitation in domesticated cotton. (A) The GLV biosynthetic pathway, initiated from linolenic acid (18:3) via 13-lipoxygenase (13-LOX), hydroperoxide lyase (HPL), (3Z):(2E) isomerase (ZEI), and alcohol dehydrogenase (ADH). (B–E) Concentrations (ng/mg) of total GLVs (B), *cis*-3-hexenal (z3HAL) (C), *trans*-2-hexenal (e2HAL) (D), and *cis*-3-hexenol (z3HOL) (E) under well-watered (WW; green symbols) and water-limited (WL; red symbols) conditions across two chemotypes, bisabolene (B) and guaiene (G), and two time points: TP1 (short-term water limitation) and TP2 (long-term water limitation). (F–G) Ratios of e2HAL:z3HAL (F) and z3HOL:z3HAL (G) used as proxies for ZEI and ADH activity, respectively. Symbols represent treatment means ± SE across 17 genotypes for the bisabolene chemotype and 6 genotypes for the Guiaene chemotype. Letters denote significant differences among groups (p < 0.05) based on Tukey-adjusted post hoc comparisons performed using the *emmeans* package in R.

In the early growing season and short-term water limitation, total GLV levels were generally consistent across chemotypes and watering regimes. However, in the later season with prolonged water-limitation, GLV production was more strongly affected by water status, particularly in the guaiene chemotype, which showed a reduction under water-limited conditions (Fig. 5B). This reduction was primarily driven by *cis*-3-hexenal (z3HAL), the most abundant GLV detected, which declined significantly in the guaiene group under prolonged drought (Fig. 5C). In contrast, the bisabolene chemotype showed a distinct short-term water-limitation response, with increased levels of the z3HAL-derived products trans-2-hexenal (e2HAL) and cis-3-hexenol (z3HOL) compared to their well-watered controls only in the early season (Fig. 5D, E).

To assess potential variation in enzyme activity across chemotypes and treatments, we calculated product-to-substrate ratios as proxies for the (3Z):(2E) isomerase (ZEI; e2HAL:z3HAL) and alcohol dehydrogenase (ADH; z3HOL:z3HAL) steps in the pathway. Both ratios were significantly elevated under short-term water limitation in the bisabolene chemotype but not in the guaiene group (Fig. 5F, G), indicating a chemotype-specific physiological response to water stress in the production of derived GLVs. By the later season, the e2HAL:z3HAL ratio increased broadly across all groups regardless of water status, suggesting a seasonal shift in isomerase activity or substrate availability (Fig. 5F). In contrast, the z3HOL:z3HAL ratio continued to show a water-limitation effect in both chemotypes (Fig. 5G), suggesting an overall enhancement in the constitutive capacity to rapidly convert z3HAL to z3HOL under drought stress.

### Genotype Exerts a Strong Impact on Terpene Concentrations

Because chemotype classifications were derived from cultivated genotypes, we examined VOC production at the genotype level to evaluate constitutive emission patterns and their modulation under water limitation. Volatile organic compound (VOC) profiles varied widely across genotypes and treatments. Monoterpene concentrations were generally higher in well-watered plants than in water-limited plants, with significant reductions observed under drought in many genotypes (Fig. 6A). These reductions were particularly evident in genotypes that exhibited the highest monoterpene levels under well-watered conditions. Chemotype effects were visible at the genotype level, with bisabolene-classified genotypes tending to produce higher monoterpene concentrations than guaiene-classified genotypes, especially under well-watered conditions. Sesquiterpene concentrations were lower overall but showed similarly high variability across genotypes, with several bisabolene chemotype genotypes exhibiting significant reductions under water-limited conditions (Fig. 6B). Notably, TM-1, the only genotype that did not produce either bisabolene or guaiene, showed a strong response to water limitation, with marked reductions in both monoterpene and sesquiterpene concentrations.

**Figure 6.**
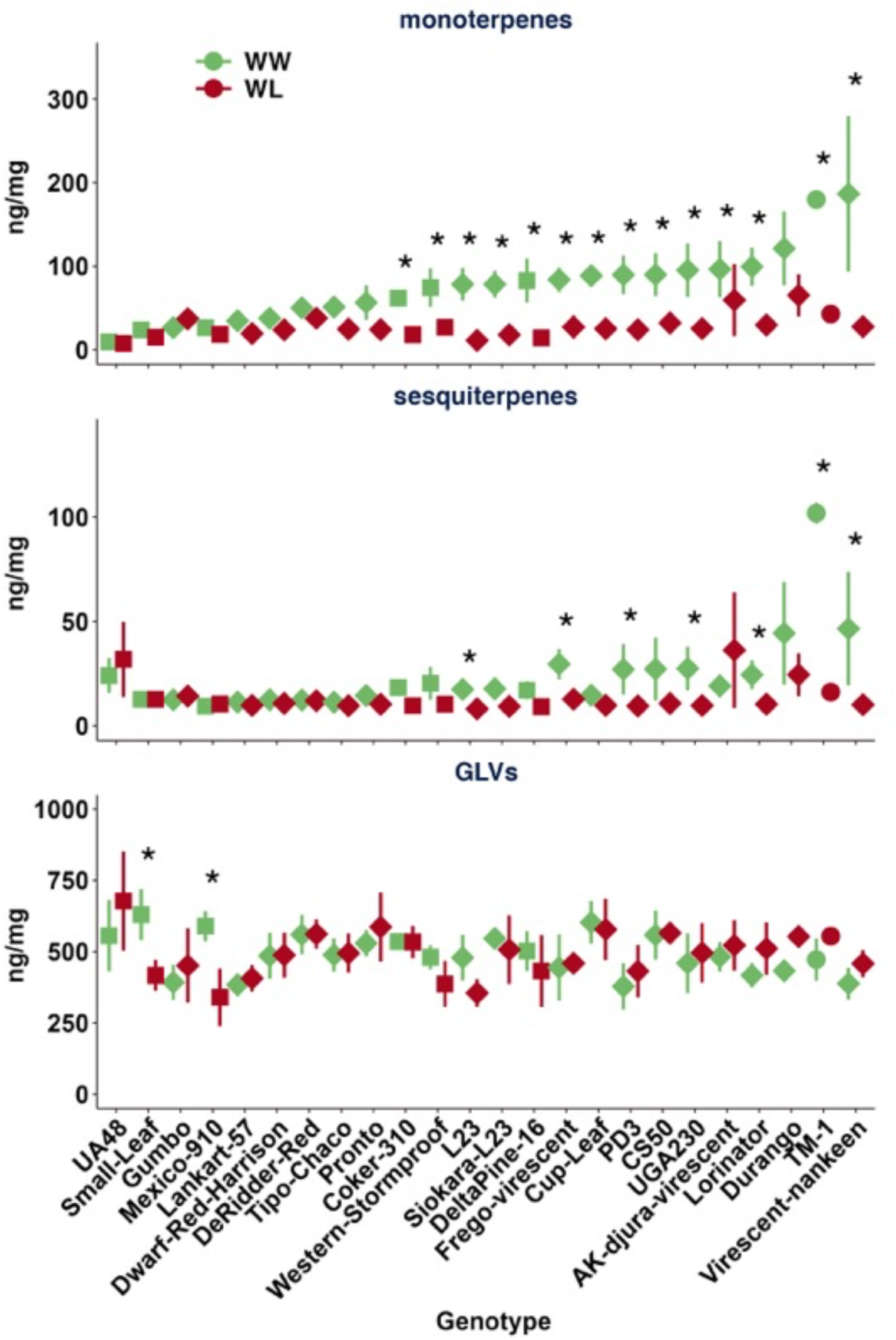
Genotypic variation in volatile organic compound (VOC) concentrations under well-watered and water-limited conditions in domesticated cotton. Mean concentrations (ng/mg) of total monoterpenes (top), total sesquiterpenes (middle), and total green leaf volatiles (GLVs; bottom) across 24 domesticated cotton genotypes under well-watered (WW; green symbols) and water-limited (WL; red symbols) conditions. Point shapes indicate chemotype: bisabolene (diamonds), guaiene (squares), and unclassified (TM-1; circle). Error bars represent ± SE. Asterisks indicate significant differences between water treatments within a genotype (*p* < 0.05) based on Tukey-adjusted pairwise comparisons using the *emmeans* package in R.

In contrast, GLV concentrations were relatively stable across water treatments for most genotypes (Fig. 5C). Only two genotypes, UA48 and Small-Leaf, showed significant differences in GLV levels between well-watered and water-limited conditions, indicating that GLV production is largely conserved under drought stress across the domesticated cotton genotypes examined.

## Discussion

We identified a previously undescribed chemotype in domesticated cotton, defined by the mutually exclusive production of either bisabolene or γ-guaiene/α-guaiene. This binary, chemotype-level distinction, present in 23 of the 24 cultivated genotypes analyzed, offers a promising opportunity to leverage VOCs as non-destructive biomarkers for assessing physiological status and stress responsiveness in cotton. As climate change accelerates the expansion of arid regions worldwide, the ability to rapidly and reliably identify genotypes with drought tolerance and resilience to complex, interacting stresses, is increasingly critical for sustaining cotton production in water-limited environments. VOCs are particularly well-suited as biomarkers in this context: they can be measured non-invasively, respond dynamically to physiological and environmental conditions, and reflect key shifts in metabolic activity. Among them, terpenes and GLVs are particularly promising due to their rapid biosynthesis, sensitivity to abiotic stress, and known roles in plant signaling and acclimation.

The chemotype-specific response in leaf water content under drought stress, with the guaiene chemotype exhibiting a more pronounced reduction compared to the bisabolene chemotype, presents a potentially key link between the chemotypic variation and physiological resilience to water limitation. As a critical indicator of plant water status, leaf water content directly impacts key physiological processes, most notably net photosynthetic rate ^54^. Water deficit triggers stomatal closure, a mechanism that conserves water by reducing transpiration^55^. However, stomatal closure concurrently restricts CO2 assimilation, impairing photosynthetic efficiency, assimilate translocation, and ultimately cotton fiber yield and quality ^56,57^. Taken together, the differential maintenance of leaf water content between these chemotypes under drought stress provides evidence that their distinct biochemical profiles are intrinsically linked to varied physiological strategies for coping with water limitation with implications for their capacity for growth and yield under adverse conditions.

The presence of distinct bisabolene- and guaiene-dominant chemotypes in cotton suggests differential regulation of specific sesquiterpene synthases. Terpene synthases (TPSs) are a diverse family of enzymes responsible for the biosynthesis of terpenes ^58^, with the TPS-a subfamily primarily associated with sesquiterpene production ^59^. In *Gossypium hirsutum*, recent genome-wide analyses have identified over 300 TPS genes, many of which are organized in clusters, indicating potential for coordinated regulation ^60^. Functional studies have demonstrated that certain TPS genes, such as GhTPS6 and GhTPS47, play roles in defense responses, with their expression being inducible under biotic stress ^58^. Notably, cotton is capable of producing both bisabolene ^31^and guaiene ^61^. While specific bisabolene and guaiene synthases have been characterized in other plant species such as Arabidopsis, Maize, Norway Spruce, Fir, Poplar, and Sunflower ^62–66^, their homologues in cotton remain to be identified and functionally validated. That said, it is known that δ-cadinene synthase can produce δ-bisabolene as a minor product when nerolidyl diphosphate is the substrate instead of farnesyl diphosphate^67^. However, this seems an unlikely source of bisabolene in our case, as δ-cadinene is ubiquitous across both chemotypes and δ-bisabolene:δ-cadinene ratios (∼1.5:1) do not align with what would be expected from δ-cadinene synthase activity alone (∼0.13:1) ^67^. Ultimately, the bisabolene/guaiene chemotypic variation may result from differences in expression levels, allelic variation, or post-transcriptional regulation of likely two distinct TPS genes. Further investigation into the expression patterns and regulatory mechanisms of candidate TPS genes in cotton could elucidate the genetic basis of this chemotypic divergence, its role in altering monoterpene biosynthesis, and its potential implications as a biomarker for stress resilience.

While chemotypic variation in wild *Gossypium hirsutum* has been documented, namely the presence of γ-terpinene-dominant and α-/β-pinene-dominant monoterpene chemotypes in populations across the Yucatán Peninsula ^36^, all genotypes analyzed in the current study fall into the low γ-terpinene group. This is consistent with the broader pattern observed in modern cultivated cotton that the high γ-terpinene group is rare or absent in breeding lines ^68^, making the discovery of a cryptic chemotypic distinction (defined by the mutually exclusive production of bisabolene or guaiene) in cultivated cotton particularly novel. This sesquiterpene-based chemotypic divergence, detected only within the domesticated gene pool, does not appear to be associated with γ-terpinene production. Its presence across nearly all modern commercial genotypes ^36^, and all the genotypes surveyed here, suggests that selection during or after domestication may have acted on alternative terpene synthase pathways, possibly due to pleiotropy or linkage with agronomic traits. These findings raise questions about how this previously undescribed chemotypic structure influences inducible defense capacity and stress responsiveness, traits with direct relevance for pest resilience and crop performance under abiotic and biotic stress.

Drought stress is known to cause substantial changes to metabolic profiles ^69,70^. Monoterpene and sesquiterpene levels reduced under water-limited conditions unmistakably indicate that their biosynthesis is sensitive to water availability, consistent with previous findings in other systems ^40,41^. This pattern was especially pronounced in cotton among genotypes with high constitutive levels of monoterpenes under well-watered conditions, indicating an active potential trade-off between growth or stress tolerance and terpene biosynthesis ^71,72^ or a biosynthetic limitation imparted by the water stress. In contrast, GLVs remained relatively stable across water treatments, with only two genotypes showing significant changes. This stability suggests that GLVs may play a more constitutive role in plant function or stress signaling in cotton that is maintained even under limited water availability. Together, these findings highlight compound class-specific responses to drought and underscore the importance of genotype-specific regulation of volatile biosynthesis under abiotic stress.

The GLV biosynthesis patterns we observed suggest that both chemotype and developmental stage shape constitutive VOC biosynthetic responses to water availability. The stability of GLV levels across genotypes early in the season, followed by drought-induced reductions in the guaiene chemotype later in the season, may indicate that this chemotype is more vulnerable to sustained water stress or employs a more conservative metabolic strategy under drought. That said, the bisabolene chemotype’s increased production of downstream GLVs (e2HAL and z3HOL) under short-term drought suggests a more dynamic or stress-responsive modulation of enzyme activity early in the season. Moreover, the shifts in e2HAL:z3HAL and z3HOL:z3HAL ratios provide indirect evidence for changes in constitutive isomerase and alcohol dehydrogenase activity, respectively. These findings support the idea that cotton can differentially regulate branches of the GLV biosynthetic pathway in response to environmental conditions, and that this regulation varies across chemotypes. Moreover, the late-season increase in z3HOL:z3HAL ratios across both chemotypes under water limitation suggests a broader, potentially adaptive shift favoring the rapid accumulation of more stable GLV products under prolonged stress ^15^. Taken together, these chemotype-specific differences in GLV pathway activity may reflect underlying physiological strategies that influence drought resilience and overall productivity.

This study focused on in vitro constitutive terpene biosynthesis and the constitutive capacity for rapid production of wound-induced GLVs, but did not assess the full spectrum of inducible volatile responses to biotic stress or plant regulated emissions in the field. Herbivory and other biotic stressors are known to induce de novo synthesis and emission of additional mono-, sesqui-, and homoterpenes ^73–76^, as well as compounds such as herbivore-inducible indole and GLV esters like *cis*-3-hexenyl acetate ^5,6,77^. These volatiles serve both direct and indirect defensive functions and mediate plant–plant signaling ^24,78–80^. Notably, we did not detect indole or *cis*-3-hexenyl acetate in our samples, likely due to the absence of herbivory from herbicide sprays applied as a standard agronomic practice at this field site. However, our findings raise important questions about whether chemotypic or genotypic variation in constitutive VOCs translates into variation in inducible responses. Given that the effect of drought on terpene biosynthesis was more pronounced in the high terpene genotypes, it is plausible that chemotypes differ not only in baseline terpene production but also in their inducibility under biotic stress. In particular, genotypes with low constitutive emissions (primarily within the guaiene chemotype) may compensate with stronger inducible responses, an idea consistent with defense optimization models ^71^. This may be particularly important in cultivated systems, where domestication has been shown to alter or reduce both constitutive and inducible volatile emissions ^23,81,82^, likely as an unintended consequence of selection for productivity and agronomic traits ^26^. These findings suggest that the history of domestication may interact with chemotypic variation to shape the capacity for inducible defense in cotton, and that evaluating inducible VOC responses may be essential for understanding the functional relevance of observed chemotype differentiation.

The one genotype that lacked both bisabolene and guaiene, TM-1, raises the possibility of a third chemotype defined not by the presence of a specific dominant compound, but by the absence of both bisabolene and guaiene. TM-1 exhibited strong constitutive production of other sesquiterpenes. In fact, TM-1 was the highest constitutive producer of sesquiterpenes across all conditions, and TM-1’s sesquiterpene production was dramatically reduced when subject to water limitation. Further investigation, including a broader survey of more diverse cotton genotypes, will be important to determine whether this represents a distinct and stable chemotype or an outlier with unique regulatory dynamics.

In summary, the discovery of a non-destructively detectable cryptic chemotype system in domesticated cotton opens the door for the development of high-throughput, non-destructive screening tools to predict genotype performance under both abiotic and biotic stress conditions. VOC signatures, particularly GLV profiles and pathway-derived ratios that are chemotype- and condition-specific, may offer novel biomarkers for breeding and selection in breeding programs aimed at developing new varieties for climate-resilient cotton systems.

## Acknowledgements

Funding for this work was provided by the BIO5 Institute (CJF), NSF IOS 2101059 (CJF), 2023310 (DP), 2102120 (DP), and Cotton Incorporated 23-926 (DP), 23-890 (DP), and 23-918 (GM).

## Contributions

C.J.F., S.J., D.P. and G.M. designed the research. D.P. and G.M. designed and performed the field experiments. C.J.F. designed the VOC sampling protocol. S.J. collected the field samples.

C.J.F. and S.J. performed the VOC collections. C.J.F. processed the data. C.J.F. and S.J. analyzed the data. C.J.F. wrote the first draft of the manuscript with contributions by all authors.

## Data Availability Statement

Raw Data from this study will be available on FigShare upon manuscript acceptance.

