## Supplemental Figures for "Identification of a volatile chemotype associated with resilience to water stress in domesticated varieties of Cotton"

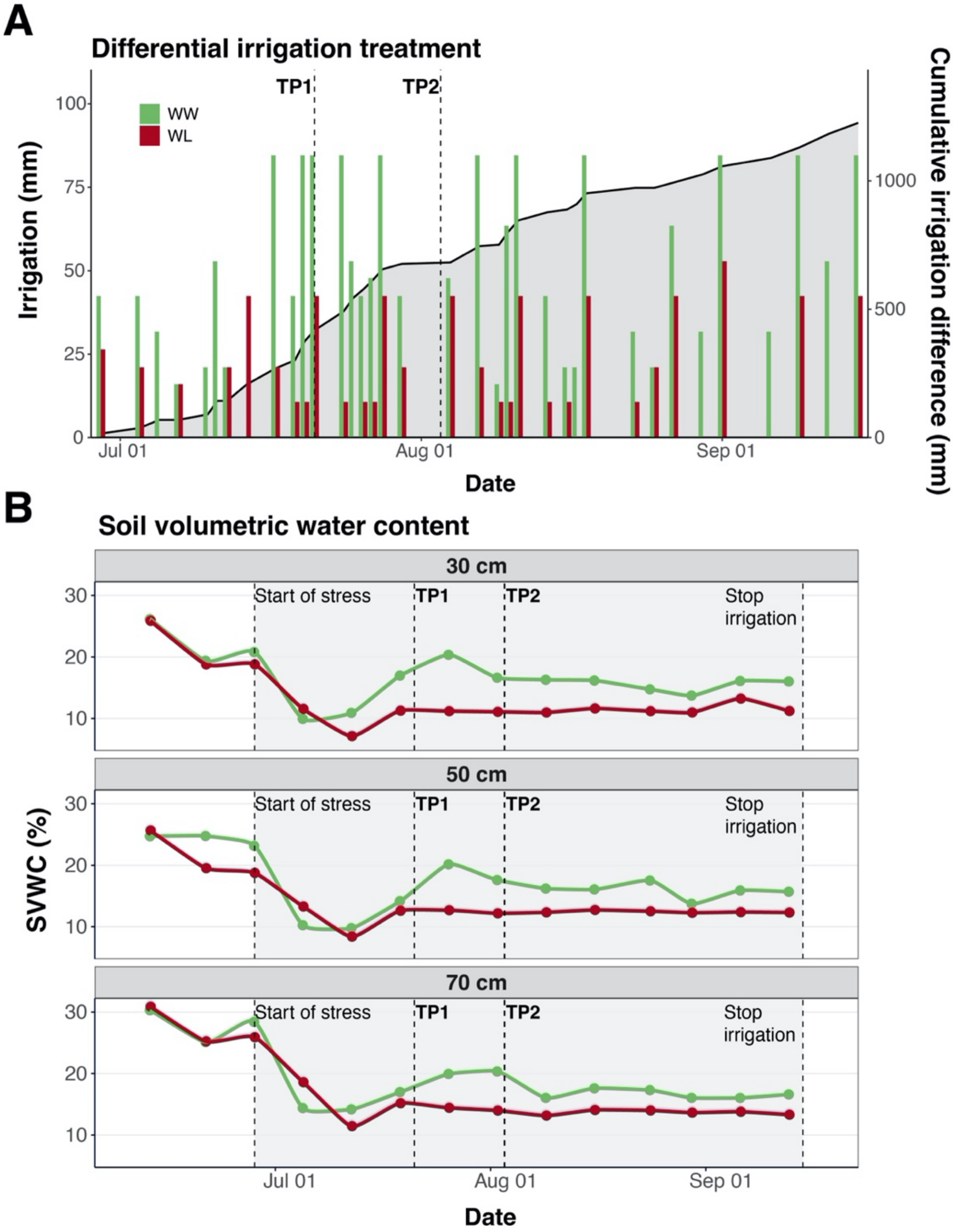


**Supplemental Figure S1. Differential irrigation regime and resulting soil volumetric water content (SVWC) profiles.** (A) Irrigation (mm) applied to well-watered (WW, green bars) and water-limited (WL, red bars) fields from July through September. The gray area illustrates the cumulative irrigation deficit in the WL treatment relative to the WW treatment. (B) Temporal dynamics of volumetric soil water content (SVWC, %) at 30, 50, and 70 cm depths for WW (green line) and WL (red line) treatments, with error bars representing standard error. The vertical dashed lines denote the initiation of the differential irrigation treatment and sampling points TP1 and TP2.

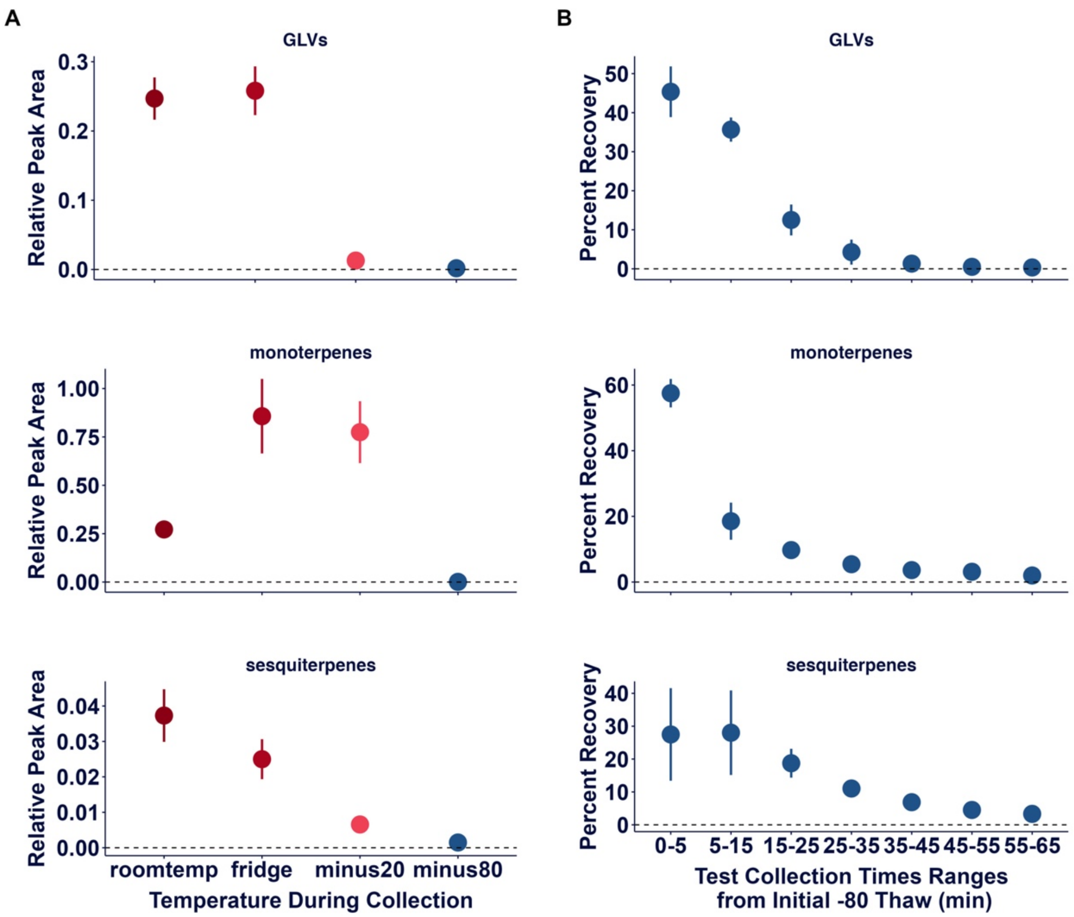


**Supplemental Figure S2. Establishing the vapor-phase VOC collection method.** (A) To confirm that storage at –80 °C prevents VOC emission from ground leaf tissue, we conducted a 60-minute collection using the flow meter apparatus described in the main text. VOCs were sampled from tubes containing frozen leaf material held at one of four temperatures: ~25 °C (room temperature), 4 °C (refrigerator), –20 °C, and –80 °C. Data represent total peak areas normalized to the internal standard (tetralin). VOC emission was negligible at –80 °C, indicating that this temperature effectively suppresses stored terpene release and GLV biosynthesis. (B) We then tested VOC emissions during thawing by moving leaf material from –80 °C to room temperature and collecting VOCs over a 65-minute period. VOCs were captured using a sequential filter system: the first filter sampled the initial 5 minutes, followed by fresh filters collecting at 10-minute intervals. This design allowed us to assess VOC recovery dynamics and optimize sampling time. Emissions declined steadily across time points, consistent with release of pre-synthesized (terpenes) or rapidly generated (GLVs) compounds. By 30 minutes, cumulative recovery reached 98% for GLVs, 91% for monoterpenes, and 85% for sesquiterpenes. By 60 minutes, cumulative recovery exceeded 99%, 98%, and 97% for GLVs, monoterpenes, and sesquiterpenes, respectively. There was no increase in VOC accumulation during the collection, indicating no detectable de novo synthesis of terpenes. Based on these results, we established 60 minutes as the standard collection duration for all assays.

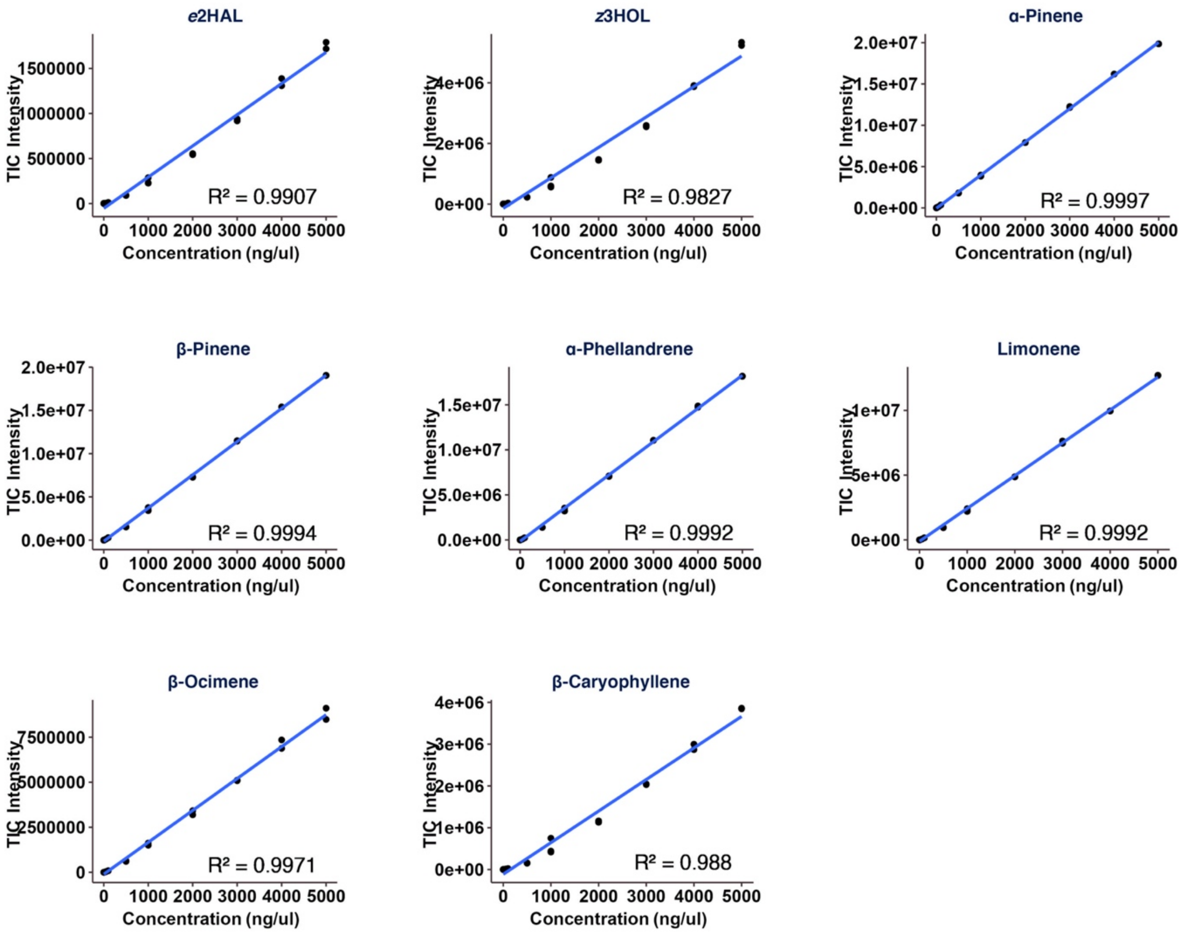


**Supplemental Figure S3. Standard curves of eight commercial standards used for quantification**. A single stock was made containing 10,000 pg/ul of each analytical standard in dichloromethane. This stock was then used to make the following concentrations: 5000, 4000, 3000, 2000, 1000, 500, 100, 50, 10, 5, 1 pg/ul. Each standard concentration was analyzed in duplicate or triplicate via GC-MS using the identical method described in the main text. Blue line represents the regression line.

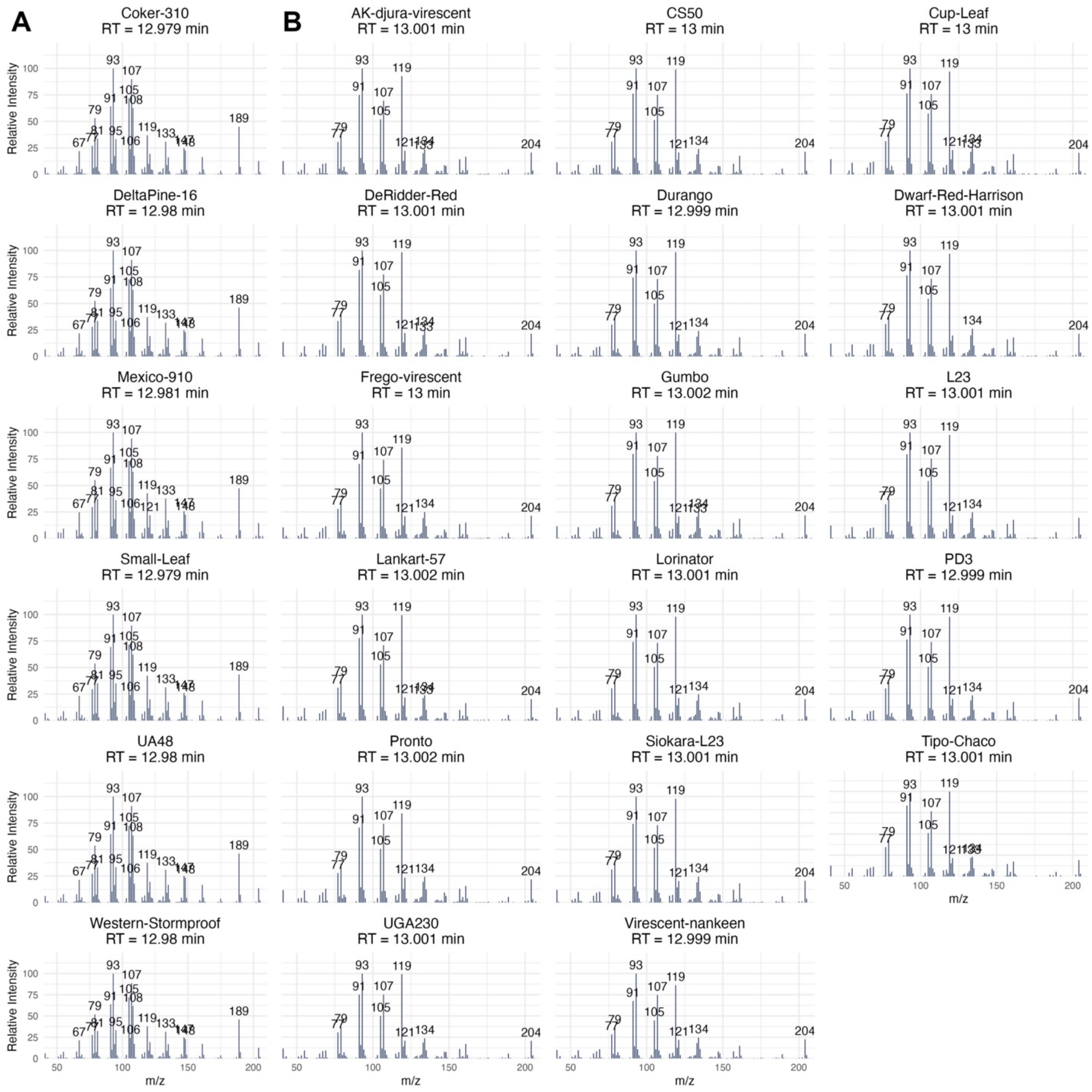


**Supplemental Figure S4. Mass Spectra of Guaiene and Bisabolene chemotypes**. The mass spectra of the compound closest to 12.99 min. (A) Six genotypes with Guiaene, showing similar mass spectra with characteristic 189 and 108 m/z, and reduced 119 m/z. Retention times are ~12.98. (B) 18 Genotypes expressing Bisabolene, with a characteristic 119 m/z, and reduced 189 and 108 m/z, and retention times ~ 13 min.
